# The secretome of a parasite alters its host’s behaviour but does not recapitulate the behavioural response to infection

**DOI:** 10.1101/799551

**Authors:** Chloé Suzanne Berger, Nadia Aubin-Horth

## Abstract

Parasites with complex life cycles have been proposed to manipulate the behaviour of their intermediate hosts to increase the probability of reaching their final host. The cause of these drastic behavioural changes could be manipulation factors released by the parasite in its environment (the secretome), but this has rarely been assessed. We studied a non-cerebral parasite, the cestode *Schistocephalus solidus*, and its intermediate host, the threespine stickleback (*Gasterosteus aculeatus*), whose response to danger becomes significantly diminished when infected. These altered behaviours appear only during late infection, when the worm is ready to reproduce in its final avian host. Sympatric host-parasite pairs show higher infection success for parasites, suggesting that the secretome effects could differ for allopatric host-parasite pairs with independent evolutionary histories. We tested the effects of secretome exposure on behaviour by using secretions from the early and late infection of *S. solidus* and by injecting them in healthy sticklebacks from a sympatric and allopatric population. Contrary to our prediction, secretome from late infection worms did not result in more risky behaviours, but secretome from early infection resulted in more cautious hosts, only in fish from the allopatric population. Our results suggest that the secretome of *Schistocephalus solidus* contains molecules that can affect host behaviour, that the causes underlying the behavioural changes in infected sticklebacks are multifactorial, and that local adaptation between host-parasite pairs may extend to the response to the parasite’s secretome content.

## INTRODUCTION

Parasites have strong impacts on the phenotype of their hosts, including modifications of the host morphology (1), physiology (2), immunity (3) and behaviour (4). Parasites with complex life cycles have been proposed to manipulate the behaviour of their hosts, in order to increase their transmission rates to their final host (5). If parasites are manipulative, they should disturb the functioning of their host central nervous system and physiological systems (hormonal and immune systems) by the production, storage, and release of molecules called manipulation factors (6-7). Parasites release a wide variety of molecules into their external environment, including nucleic acids (8) and proteins (9), which are sometimes included in extracellular vesicles (10). These molecules are collectively referred to as the secretome. While the effects of exposing hosts to the secretome of their parasite on their immune response has been demonstrated in some host-parasite pairs (11, 12), there is currently no experimental evidence that the secretome content from a parasite directly affects behaviour in the host in the same way that the whole parasite infection does (13).

We studied the effect of the parasite secretome on host behaviour using the interaction between the threespine stickleback and the parasite *Schistocephalus solidus. Schistocephalus solidus* is a freshwater cestode that is trophically transmitted to three hosts (14). It enters the body cavity of its first host, a copepod, after being ingested, and then negatively affects its growth and reproduction (15). In the copepod, the expression of risky behaviours shows a pattern of “protection then facilitation” (16-17): infected copepods exhibit reduced activity in the early phase of infection, then increased activity and a changed swimming pattern, leading to higher detection and ingestion by the second host, the threespine stickleback (18). Infection in this host results in changes in morphology, physiology, immune response and behaviour (19). The worm life cycle is completed when it reaches its final avian host. (19).

The parasite’s life stage within the body cavity of the threespine stickleback is characterised by two phases with distinct phenotypes in both the parasite and its host. During the early days of infection (the non-infective phase), the worm shows high growth and is too small to mature if switched to its final bird host (20). The host does not show the expected adaptive immune response (21) and no behavioural changes are known (22). At the infective phase, the worm is ready to reproduce in its final avian host (20). The transition from the non-infective to the infective phase is characterized by major genome-wide reprogramming events in the worm, including the activation of genes predicted to be involved in neural pathways and sensory perception (23). In this infective phase, drastic behavioural changes that result in a loss of the anti-predator response appear in sticklebacks (22). Compared to non-infected individuals, they are more exploratory (24), less anxious (25) bolder in the presence of a predator (26), they recover more quickly from a frightening overhead stimulus, and feed following the stimulus (26). These behavioural perturbations could result in part from the combined effects of modifying the serotonin pathway and the immune response in infected fish (21, 27, 28). At the infective phase, secretome collected from *S. solidus* has been shown to modulate the stickleback immune response *in vitro* (29). Whether this modulation of the immune system is also involved in behavioural changes is unclear (27, 28).

Altogether, the timing and the nature of these changes in both host and parasite support the hypothesis of a manipulation of the threespine stickleback by *Schistocephalus solidus* to reach its final avian host. The tegument of *S. solidus* is known to include vesicles (30). However, whether the worm secretes any molecule that can be classified as manipulation factors that are able to alter fish behaviour is unknown (7). This system offers a unique opportunity to functionally test if the secretome is responsible for the behavioural changes. We predict that only exposure to secretome from worms at the late, infective phase should induce behavioural perturbations similar to what is seen in infected sticklebacks, and not exposure to secretome from worms at the early, non-infective phase.

The interaction between *S. solidus* and the threespine stickleback is highly specific, since it is the only fish host that this parasite can infect (31). *S. solidus* infects freshwater sticklebacks (19) and a combination of studies of wild populations and of experimental reciprocal cross infection data suggests local adaptation between sympatric (“native”) host-parasite pairs (32-34). On the other hand, marine sticklebacks are rarely exposed to *S. solidus*, as the worm does not tolerate high salinity levels when its hatches (35). Potentially reflecting the lower probability of co-evolution between these allopatric (“naïve”) marine fish and *S. solidus*, infection success of experimental infections is highly variable, ranging from 1.4% to 99% depending on the population (34, 36). Marine fish infected by *S. solidus* show morphological changes (37) and the immune response of allopatric fish exposed *in vitro* to *S. solidus* antigens at the infective phase differs from the one of sympatric sticklebacks (38). However, the effects on behaviour of infecting an allopatric stickleback with *S. solidus* have never been studied.

The occurrence of threespine stickleback populations from freshwater and marine environments and their variation in infection susceptibility allow us to functionally test if *S. solidus* and its secretome co-evolved with its sympatric freshwater host to specifically modify its behaviour. Strong co-evolution between a manipulative parasite and its sympatric host was previously reported in the *Ophiocordyceps*-ant system, where *ex vivo* essays demonstrated that the fungus *Ophiocordyceps* secretes a specific array of metabolites only in presence of the host brain it has specifically evolved to manipulate (39). We had two opposite but plausible predictions according to the manipulation hypothesis: the first one was based on the observation that marine fish can be highly susceptible to *S. solidus* infection (34). If allopatric sticklebacks do not have adaptive mechanisms to fight against the secretome effects because they did not evolve with the parasite, we predicted that they would be more sensitive to the secretome than the sympatric population and would show stronger behavioural changes. The second prediction was that marine fish would activate their immune response earlier than sympatric fish following infection, which would reduce the effects of secretome on fish physiology and behaviour (40). In this case, we predicted that the allopatric population would be less sensitive to the secretome from the infective phase than sympatric fish and that they would show reduced or no behavioural changes. In both cases, we predicted that the injection of secretome from the non-infective phase would not modify behaviour of allopatric fish.

The objective of this study was to test if the secretome of a parasite located outside the brain is sufficient to induce behavioural changes in its vertebrate host. To reach this objective, we used a functional approach (Figure 1).

**Figure 1.**
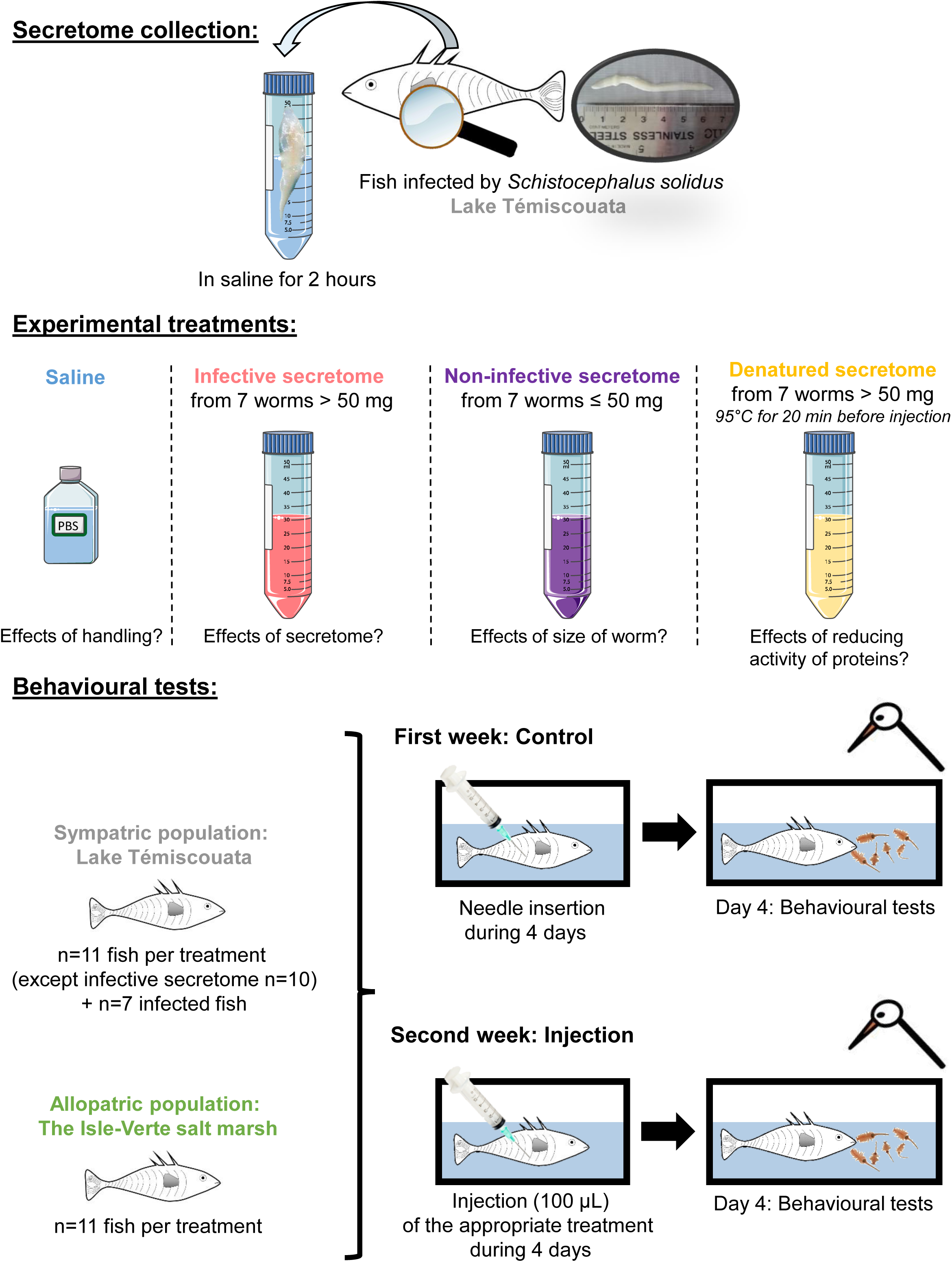
Experimental design used to determine if the secretome of a parasite is sufficient to induce behavioural changes in its vertebrate host. Secretomes from the non-infective and infective phases of *S. solidus* were injected in non-infected sticklebacks from a sympatric and an allopatric population. Only injection of the infective secretome was predicted to induce behavioural changes. Saline (PBS), non-infective secretome, and denatured secretome treatments were used as controls for which no behavioural changes were expected. In the sympatric population, infected fish were also tested to validate that the behavioural tests could discriminate between healthy and *S. solidus* infected fish.

## MATERIALS AND METHODS

### Collecting secretome from the non-infective and infective phases

The collection of the secretome from *S. solidus* was performed according to a protocol adapted from (41). We caught threespine stickleback fish from Lake Témiscouata (Québec, 47°80’ N 68°87’ O), in June and July 2016 using minnow traps. One day after fishing, fish (n=41) were euthanized with an overdose of MS-222 (75 mg/L mg/kg) and dissected to determine if they were infected by *S. solidus*. Fish sex, size and mass, and *S. solidus* mass and number in each fish were noted. Twenty-eight fish were infected by worms, 8 at the non-infective phase (worm mass ≤ 50 mg) and 20 at the infective phase (worm mass > 50 mg;). Following dissection, each *S. solidus* worm was washed to remove fish blood and placed in a saline solution protected from light to collect its secretome for 2 hours (see Supplementary material). The secretome was stored at -20°C. For subsequent injections, we only used secretomes sampled from fish harbouring a single worm (n=21, including 7 non-infective secretomes and 14 infective secretomes) in order to remove potential effects of multiple infection on secretome content and function.

### Sampling of sympatric and allopatric fish hosts

We performed the injections in two populations of threespine sticklebacks. The first population, from Lake Témiscouata, Québec, was composed of fish that lived in sympatry with the worms used to obtain secretome samples. The second population was composed of fish from a salt march at Isle-Verte, Québec (48°00’ N 69°40’ O; salinity 22-26 ppt), which thus lived in allopatry with the worms used to obtain the secretome. Fish in this population are anadromous, meaning that they live in saltwater and migrate to the lower salinities of the salt marsh to reproduce. Sticklebacks in this population are not expected to be infected by *S. solidus*, because the worm does not tolerate high salinity (42). Detailed sampling in Supplementary material.

### Experimental treatments

We used four treatments for each population: an injection control treatment, an “infective secretome” treatment, a “non-infective secretome” treatment and a “denatured infective secretome” treatment. For these four treatments, we used healthy non-infected fish from the two populations. In addition, in the Témiscouata population, we added a group of fish infected by *S. solidus* that were not injected as a positive control, to validate that the behavioural test could discriminate between healthy and *S. solidus* infected fish (n=7). The control treatment involved fish injected with phosphate-buffered saline (PBS), to test handling effects on fish behaviour. The three other treatments involved injections of secretome collected in Lake Témiscouata. For each of these three treatments, we prepared a mixture of secretome that insured sufficient volume for injections, for a total of 3 mixtures (“infective secretome”, “non-infective secretome” and “denatured infective secretome”). For each treatment, we used one secretome mixture from 7 worms to remove potential individual effects from a specific worm: if a worm secreted proteins with aberrant effects on fish behaviour, then their effects would be more likely to be buffered by the proteins secreted by the other six worms. For each treatment, the same mixture was injected in fish from the Témiscouata and the Isle-verte populations, which allowed to compare behavioural effects between them. Each mixture was made of 1600 µL of secretome from each worm. The infective secretome treatment included fish injected with secretome from worms whose weight was above 50 mg (the mass at which the worm becomes infective (19)). Mass of the worms ranged from 80 mg to 700 mg with a mean of 368.6 mg. The non-infective secretome treatment used fish injected with secretome from worms whose weight was equal to or under 50 mg. Worms had a mass around 10 mg (minimal value detected by the balance) to 50 mg with a mean of 27.1 mg. This treatment was used as a control for which behavioural changes were not expected. The denatured secretome treatment used fish injected with secretome from worms whose weight was above 50 mg, but for which the proteins included in the secretome were denatured by heat at 95°C for 20 minutes before each injection. This treatment was also added as a negative control, as proteins lose their secondary and tertiary structures following heat treatment, and consequently their biological activities. If we found differences in behaviours between fish injected with the intact and the denatured secretome from large worms, this would suggest that the activity of proteins included in the secretome is involved in the behavioural alteration. Worm mass for this last treatment was comprised between 170 mg and 730 mg with a mean of 464.3 mg. Each treatment was composed of n=11 fish in each population, except for the infective secretome treatment in the Témiscouata population (n=10). We verified protein concentration and content of the secretome mixtures using a Bradford assay and SDS-Page (Supplementary material, supp. Fig 1).

### Injection protocol

At the beginning of the protocol, we isolated 51 fish from the Témiscouata population and 44 fish from the Isle-Verte population in 2-L tanks and assigned an identification number (ID). The first day following isolation, we verified infection by *S. solidus* using the environmental DNA method (43). Forty-four fish from the Témiscouata population were detected as non-infected and could be used for injections, while 7 fish were infected by *S. solidus* and were used as control infected fish. As expected, none of the fish from the allopatric population was infected by *S. solidus* and all could be used for injections.

We randomly assigned each fish to an injection treatment (PBS, “infective secretome”, “non-infective secretome” or “denatured infective secretome”). During the first week (control week), we inserted a needle into the abdominal cavity of each fish (no liquid injected), once a day for four days. On the 4^th^ day, we tested behaviour of each fish between 30 and 45 minutes after manipulation. During the second week (injection week), we injected each fish in the abdominal cavity with 100 µL of the appropriate treatment, once a day for four days (total of 400 µL injected per fish). On the 4^th^ day, we tested behaviour of each fish between 30 and 45 minutes after the last injection. Denatured secretome was heated at 95°C for 20 minutes and was cooled before each injection. Infected fish used as a positive control were exposed to the same protocol, except that during the injection week the needle was only inserted. Fish were not anesthetized, as manipulation took less than 1 minute and induced minor stress to fish. We fed fish daily with brine shrimps following needle insertion or injection, except on day 4. Each fish was given 3 days of rest between the control and the injection weeks. One fish for the infective secretome treatment of the Témiscouata population died between the two weeks and was removed from analysis.

### Behavioural tests and analysis

The behaviours of each fish were measured in a 45 L test aquarium using standard tests to measure exploration, thigmotaxis and boldness in threespine sticklebacks (22, 25, 27) (see Supplementary material). Statistical analyses were performed using the R software version 3.5.0 (44). See supplementary material for additional information on statistical tests.

## RESULTS

### Behaviours that differentiated infected and non-infected fish

We measured seven behaviours in order to quantify exploration, thigmotaxis (wall-hugging tendency (25)) and boldness in infected and non-infected fish. We found that in the sympatric population, infected and non-infected fish differed significantly in two behaviours associated with boldness: latency to feed before a predator attack and time spent frozen after a predator attack. Infected fish ate two times more rapidly before a predator attack (average measured in control week: 113 sec) than non-infected fish (average measured in control week: 224 sec) (p = 0.049) (Figure 2 panel 1.A). Infected fish almost did not freeze after the predator attack (average measured in control week: 6 sec) while non-infected fish froze for a longer time (average measured in control week: 107 sec) (p = 0.009) (Figure 2 panel 1.B). For the other five behaviours, infected fish did not differ significantly from non-infected fish, although trends were apparent (Figure 2, panel 2).

**Figure 2.**
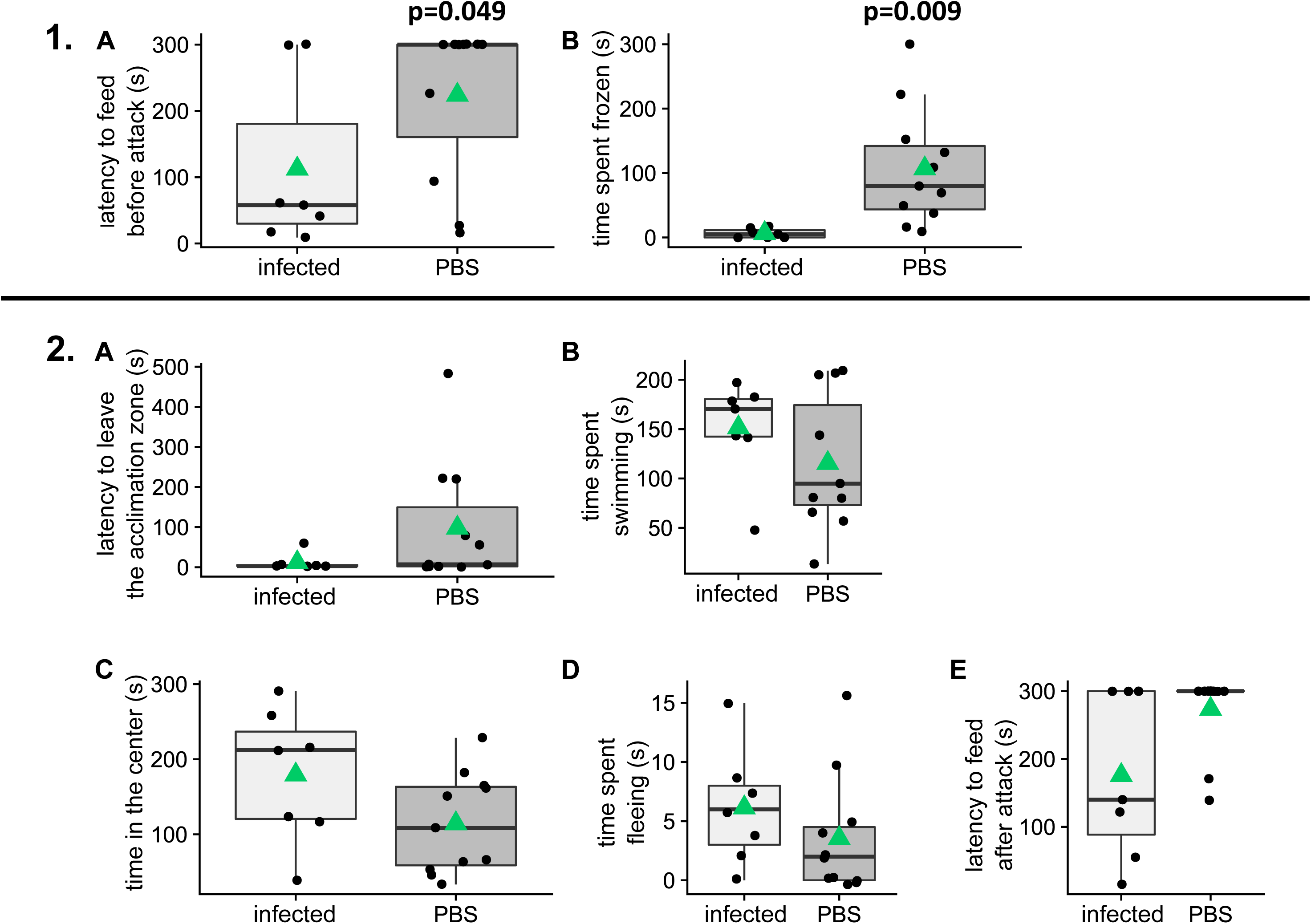
Two boldness-related behaviours differ between *S. solidus*-infected and non-infected sticklebacks. **1.** Behaviours that were significantly different between infected and non-infected sticklebacks (p < 0.05). A. latency to feed before a predator attack (sec). B. time spent frozen after a predator attack (sec). **2.** Behaviours that were non-significantly different between infected and non-infected sticklebacks. A. latency to leave the acclimation zone (sec). B. time spent swimming (sec). C. time in the centre (sec). D. time spent fleeing after a predator attack (sec). E. latency to feed after a predator attack (sec). Each comparison was performed between infected fish and non-infected fish (from the saline (PBS) group) of the control week. Black lines represent the median, the green triangles the average, and each dot one individual.

### Behavioural effects of secretome injections in non-infected fish from two populations of threespine sticklebacks

#### Sympatric population: Lake Témiscouata

We tested behaviour twice in each individual: during the control week and during the injection week. We first tested control non-infected individuals in which we injected only a saline solution, in order to quantify any changes related to the disturbance of the experiment in these two weeks. For all behaviours assessed, we did not find any significant differences between the two weeks in these control fish, suggesting that handling and injections did not significantly affect them. Similarly, infected fish did not change their behaviour between the first and second tests (Supp Tables 1 to 7).

We were not able to reproduce the typical behaviours of infected fish using secretome injections from the infective phase when injecting healthy individuals from the Témiscouata freshwater population (Supp Tables 1 to 7, Supp Figures 2 to 7). For example, in this sympatric population, none of the secretome injections had significant effects on feeding latency before a predator attack, while it did affect fish from the allopatric population (see below). Sympatric fish injected with the infective secretome did not significantly change the time they took before eating between the two trials (p = 0.975, Supp Table 4, figure 3 panel 1.C). The same was observed in fish injected with the non-infective secretome (p = 0.852, Supp Table 4, figure 3 panel 1.D) and following injection of denatured secretome (p = 0.526, Supp Table 4, figure 3 panel 1.E).

**Figure 3.**
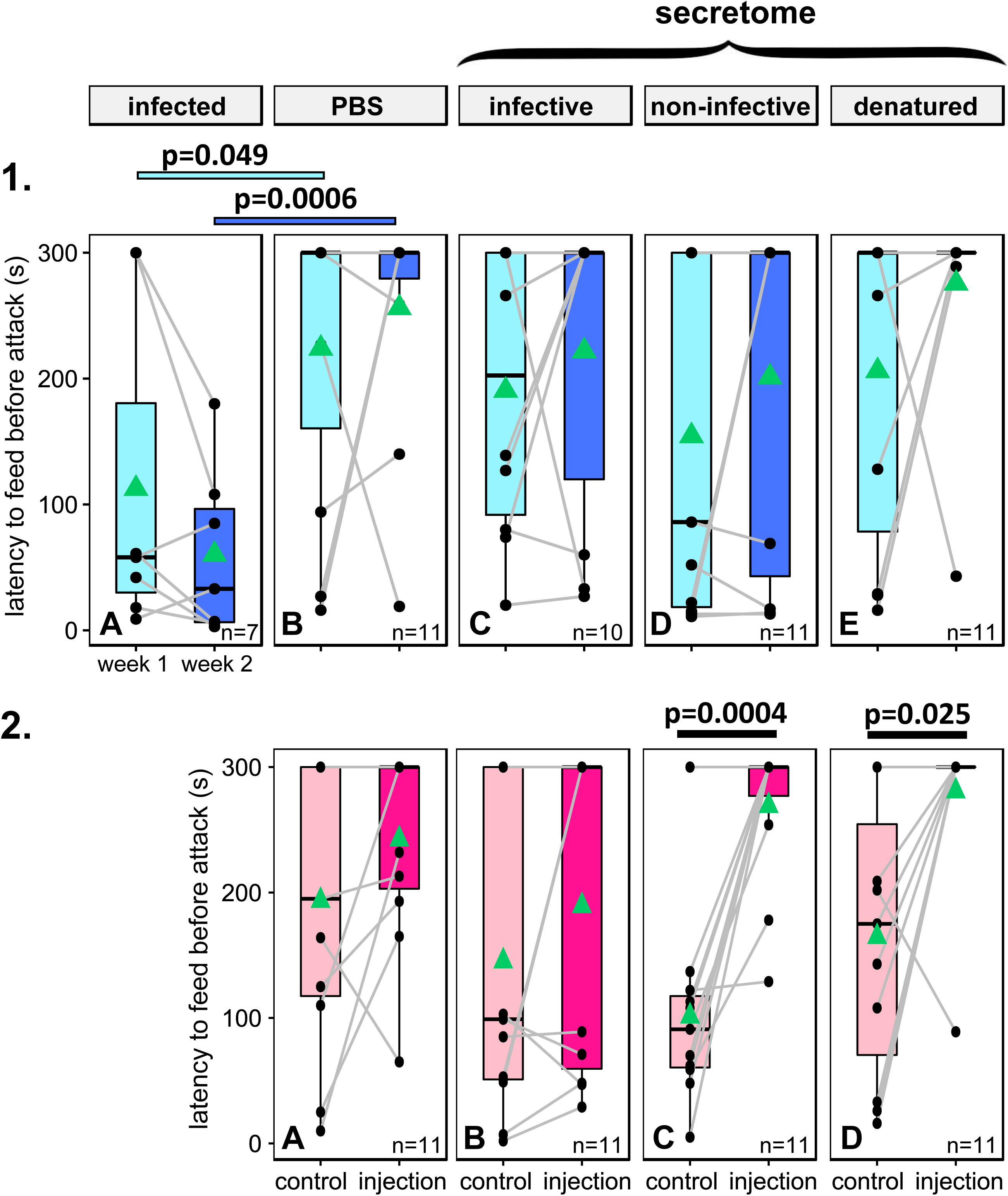
Injections of non-infective secretome significantly lower the expression of boldness-related behaviours, only in fish from the allopatric population. **1.** Latency to feed before a predator attack (sec) in the Témiscouata population. A. Infected fish. B. Fish injected with control saline (PBS). C. Fish injected with infective secretome. D. Fish injected with non-infective secretome. E. Fish injected with denatured secretome. **2.** Latency to feed before a predator attack (sec) in the allopatric population. A. Fish injected with control saline (PBS). B. Fish injected with infective secretome. C. Fish injected with non-infective secretome. D. Fish injected with denatured secretome. Black lines represent the median, the green triangles the average, each dot one individual, and grey lines the reaction norm of one individual between the control and injection weeks.

#### Allopatric population: The Isle-Verte salt marsh

In the allopatric population, we first tested control individuals in which we injected only a saline solution, in order to quantify any changes related to the disturbance of the experiment in these two trials. For all behaviours assessed, we did not find any significant differences between the two weeks in these control fish, suggesting that handling and injections did not significantly affect them (Supp Tables 8 to 14). For example, the control non-infected fish had similar latency to eat before the predator attack in the first and second weeks (p = 0.762, Supp Table 11) (Figure 3 panel 2.A).

We found that behaviour was changed after secretome injections in the allopatric population in some treatments. The latency to eat before a predator attack was not changed after injection of the infective secretome in healthy individuals (p = 0.642, Supp Table 11, figure 3 panel 2.B). However, fish injected with the non-infective secretome significantly increased their latency to approach food, taking more than twice longer before resuming feeding (p < 0.001, Supp Table 11, figure 3 panel 2.C). After injection of denatured secretome, fish significantly delayed or refrained entirely from eating before the predator attack compared to the control week (p=0.025, Supp Table 11, figure 3 panel 2.D). We also found that allopatric fish took significantly longer to resume feeding after a predator attack following injection of the non-infective secretome (p=0.037, Supp Table 14). After a predator attack, they stopped feeding entirely during the injection week (Supp Figure 7 panel 2.C). This behaviour was not significantly affected in any other treatments (Supp Table 14, supp figure 7 panel 2). Additionally, we found that allopatric fish significantly spent more time swimming after injection of denatured secretome (p=0.016, Supp Table 9), while other secretome treatments did not affect this behaviour (Supp Figure 3 panel 2.D). None of the other behaviours showed changes after injections of secretome in this population (Supp Tables 8, 10, 12 and 13 and Supp Figures 2 to 7).

## DISCUSSION

It has been proposed that parasites manipulate the behaviour of their host by secreting molecules (*i.e.* manipulation factors) that ultimately impact the physiology and the behaviour of their host. Here, we tested if the secreted molecules of *S. solidus* could be responsible (entirely or in part) for the behavioural changes in infected sticklebacks, by injecting secretome in fish from a native, sympatric population and from a naïve, allopatric one. We were not able to recreate the typical risky behaviours of infected fish in healthy individuals using solely the secretome of *S. solidus* from the infective phase. However, the secretome of the parasite was sufficient to significantly alter two risk-taking behaviours, the latency to feed before and after a predator attack, in fish injected with the secretome from early infection worms, but only in the allopatric fish population that did not evolve in presence of the parasite.

The lack of behavioural change in fish injected with infective secretome can be interpreted in two ways: either the secretome is not sufficient to alter behaviour, or our protocol did not recreate the biological alterations that happen in the host during the weeks of infection. In both populations, fish injected with the infective secretome did not differ in behaviour from the control non-infected fish, despite the fact that proteins were detected in the secretome. It is possible that *S. solidus* does not act on the behaviour of its host through its secretome. If such is the case, the probability that this host behaviour alteration is the result of active manipulation by the parasite would be low. If this is the correct interpretation, behavioural changes would be the result of a side effect of infection, such as the parasite mass burden, or the result of a host response (25). However, it would be improbable to observe the significant changes in behaviour we quantified when injecting the non-infective secretome. Moreover, as our behavioural assay could detect significant differences in latency to feed before a predator attack between infected and non-infected fish, we expect that we also had the statistical power to detect differences in the infective secretome treatment.

An alternative explanation would be that our exposure protocol was not sufficient to test our prediction. First, it is possible that we did not reach a threshold level of manipulation factors, as it has been proposed that they may be accumulated slowly and stored by the parasite (7). Because secretomes were sampled from the wild, the amount of secretome collected was limited by the number of fish caught that were parasitized by infective *S. solidus*. We thus injected a small dose every 24 hours for 4 days. In contrast, if manipulative, *S. solidus* probably secretes molecules continuously within its host. Increasing the dose and exposure duration should definitely be tested in order to better understand the role, if any, of the parasitic secretome in host manipulation. Second, a potential explanation is that *S. solidus* may secrete molecules during the non-infective phase that would act to “prime” the host (13). One could test the importance of the interaction between the effects of molecules secreted in the non-infective and infective phases in inducing behavioural changes by sequentially injecting a fish with the non-infective and infective secretomes. Third, our protocol did not include the potential combined effects on behaviour of the secretome and of the physical deformations induced by the worm in the fish abdominal cavity. This could be tested by simultaneously injecting the infective secretome with silicone, which was previously used to re-create the parasitic mass burden in non-infected fish (25). Finally, it is also possible that the infective secretome acts in combination with molecules of the stress response, which was previously proposed to be involved in the host’s behavioural changes (27, 28). A next step will require to adopt an integrative functional approach to simultaneously test the effects of the secretome, of the activation of the stress response, and of the parasitic mass burden on host behaviour.

The significant increase in latency to eat before and after a predator attack induced by the secretome from worms at the non-infective phase is similar to the altered behaviour seen in infected copepods, the first intermediate host of *S. solidus.* The copepod host exhibits a behaviour pattern of “protection then facilitation”, first showing reduced activity in the early phase of infection, presumably reducing detection by predators, then an increased activity, which leads to higher detection by threespine sticklebacks (16-18). The induced behavioural perturbations in sticklebacks are surprising, as behavioural changes have not been reported in infected sticklebacks when *S. solidus* is in the early non-infective phase (22). It is possible that the same shift in behaviour exists in the fish host but has gone undetected. Using single experimental infection in 5 female sticklebacks, Barber et al. tracked changes in their antipredator behaviour for 16 weeks post-parasite exposure and reported significant behaviour changes only when the worm reached 50 mg (22). This could be explained in part by their low sample size, and testing more fish infected by worms at the non-infective stage could provide a better understanding of the potential behavioural sequence of “protection then facilitation”. We can hypothesize that the secretome contributes to the worm protection at the non-infective stage, while other mechanisms (including the host stress response and the parasitic mass burden) could act independently or in combination with the infective secretome to alter behaviours that facilitate worm transfer to the final host. Also, it is important to keep in mind that secretomes were sampled from naturally infected fish, which prevented to precisely estimate the developmental stage of each worm. For example, small worms (≤ 50 mg), which we considered non-infective, might have grown more slowly because they invested more resources into molecules responsible for behavioural changes. Finally, the divergence in behavioural effects between the non-infective and the infective secretomes suggests that the secretome content between these two phases is different. In accordance with our results, the analysis of the transcriptome of *S. solidus* previously demonstrated that the genes expressed by the worm in the non-infective and infective phases were associated with contrasting biological functions, with thousands of genes differing in activity (23). However, it is also possible that behavioural differences result solely from concentration effects, if non-infective worms produced less concentrated secretomes compared to infective worms when their secretions were collected in a saline solution of constant volume. Analysing the nature of the protein content of the *S. solidus* secretome at each stage would provide functional information that could shed light on the nature of the molecular interaction between the threespine stickleback and *S. solidus*.

Studies in several host-parasite pairs from different branches of the tree of life have shown that behavioural alteration in the host affects specific behaviours rather than disrupting the host response to all cues (5, 7, 19, 45). Not all behaviours were altered by the non-infective secretome in our study. The specificity of effects on cautiousness in presence of a predator, but not on exploration of a novel and risky environment, or of anxiety-like behaviour such as wall hugging, is in accordance with previous studies that have shown that not all behaviours are modified in infected sticklebacks (45) and that not all behaviours are changed when healthy sticklebacks are experimentally manipulated to behave like infected ones (27). Indeed, manipulating serotonin levels in healthy sticklebacks to mimic the changes observed in the brain of infected ones lead to changes in schooling and the tendency to swim close to the surface, but not the response to a predator attack, which was significantly altered in the opposite direction to an infected fish in the present study (27). This result points to a probable multifactorial nature of host altered behaviours, as no experimental manipulation leads to changes in all the behaviours observed in *Schistocephalus* infected fish. Here, the non-infective secretome exposure resulted in more cautious fish that took longer to feed under risk. Identifying the content of the secretome will potentially uncover molecules that affect feeding directly or indirectly, as found in another host-parasite system where feeding is affected (13).

Our results show that the secretome of *S. solidus* had significant effects on fish risky behaviour only in an allopatric (naïve) population. This result is in accordance with the hypothesis that since allopatric sticklebacks did not evolve with *S. solidus*, they do not possess the adaptive “generalised response” against *S. solidus* and its effects on fish physiology and behaviour, as proposed for this parasite and others (34, 46). Local adaptation and potential co-evolution between the threespine stickleback and *S. solidus* was previously suggested with cross-infection studies (33, 43, 63). Our results point to the need for further studies on local adaptation in this host-parasite pair, by including secretome exposure of fish that have evolved in the presence of another worm genotype, *i.e.* sticklebacks of freshwater origin that can recognize the parasite but did not co-evolve specifically with that genotype (34, 47). For instance, non-infected freshwater sticklebacks from the west Coast of Canada (known to be infected by *S. solidus* (34)) could be exposed to the secretome from Lake Temiscouata worms.

The significant increase in the latency to eat before an attack, which was reported after the injection of the non-infective secretome in the allopatric population, was also found in fish injected with heat-denatured secretome, suggesting that these impacts on behaviour were not related solely to the activity of proteins. Indeed, since these secretome samples were denatured by heat in order to lose the secondary and tertiary structures of proteins, and consequently their biological activities, it is possible that this behavioural change might be induced by other molecules included in the secretome that were not impacted by the heat treatment. It is likely that *Schistocephalus* secretes and/or excretes other molecules, such as extracellular vesicles including micro-RNAs (11). These molecules could have their biological activities reduced in the presence of the active secretome proteins, and heat treatment possibly revealed their actions. To confirm this hypothesis in the future, we could replace the heat treatment of secretome by a protease exposure, in order to specifically break down and inactivate proteins in the secretome. Heat treatment and protease exposure could also be applied to the non-infective secretome to better understand the role of proteins and other molecules in the reported behavioural change.

In this study, secretome alone did not recapitulate the behavioural changes of infected sticklebacks in non-infected sticklebacks. Yet, the effects of secretome from the non-infective phase on boldness in the allopatric population suggest that there may be a “protection then facilitation” effect at play in the threespine stickleback host, as seen in the first copepod host. Our findings demonstrate that the secretome of *S. solidus* is an important component of the molecular cross-talk between the parasite and its threespine stickleback host.

## Supporting information

Complete dataset

## Acknowledgements

The authors thank the personnel of the Laboratoire Aquatique de Recherche en Sciences Environnementales et Médicales at Université Laval for their help with fish rearing. We thank Christian Landry for his comments and advice during the development of this project. We thank Marie-Pier Brochu, Verônica Alves, Florent Sylvestre and Sann Delaive for comments on previous versions of this manuscript. This project was funded by an NSERC Discovery grant to NAH. The authors declare no competing or financial interests.

## Author contributions

C.S.B. designed the study with input from N.A.H.. C.S.B. performed the experiments. C.S.B. analysed the data with input from N.A.H.. C.S.B. and N.AH. wrote the manuscript.

## Ethics

This study was approved by the Comité de Protection des Animaux de l’Université Laval (CPAUL 2014069-3).

## Data availability

The complete dataset is available as a supplementary tab-delimited .txt file. Data on seven behaviours measured during the control and treatment weeks, mass (g), size (mm) and sex of each fish are reported, as well as the infection status (no=0; yes=1), the number and the total mass of *S. solidus* (g) for individual fish in each treatment.

## Electronic supplementary material

A) Supplementary tables. Results of all the statistical comparisons performed with the lmer function to determine if the secretome injections change the behaviour of non-infected fish of the Témiscouata population and of the allopatric population are presented. Results are reported in separate tables for each of the seven behaviours and for each population (supplementary tables 1 to 14). B) Supplementary figures. Seven supplementary figures are presented. C) Supplementary information on methods.

