## Supplementary figures and images for "The secretome of a parasite alters its host’s behaviour but does not recapitulate the behavioural response to infection"

### Supplementary figures

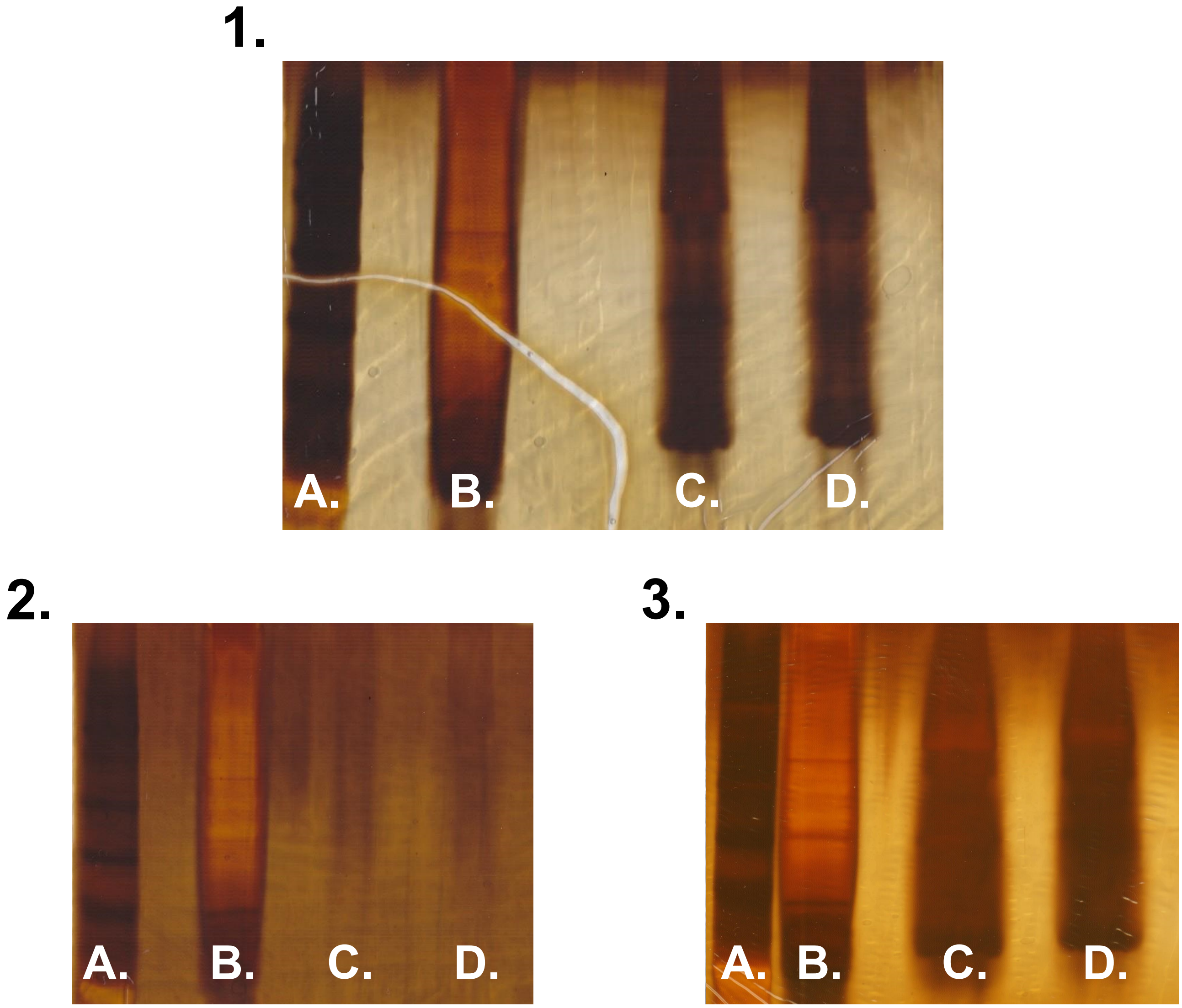

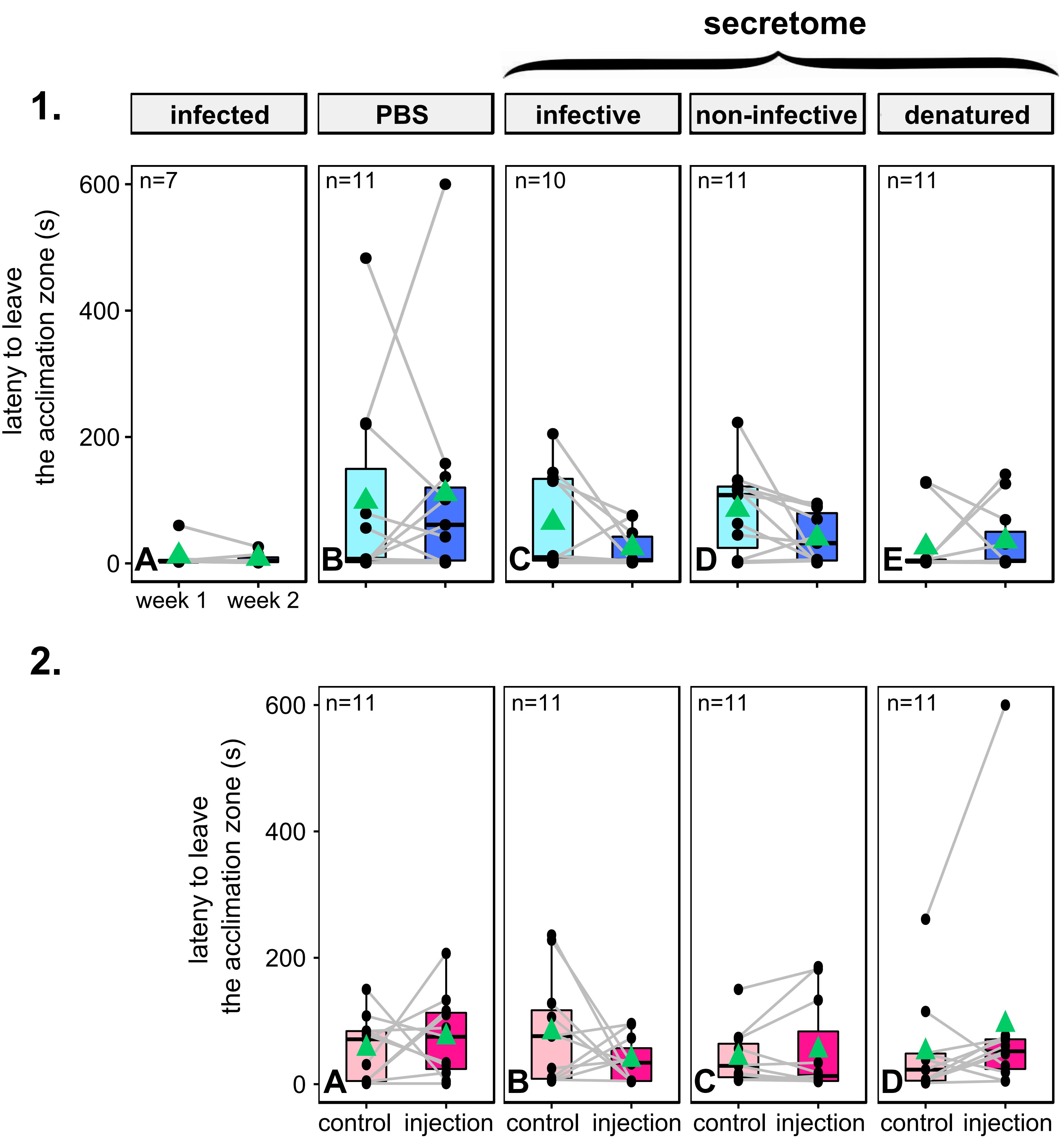

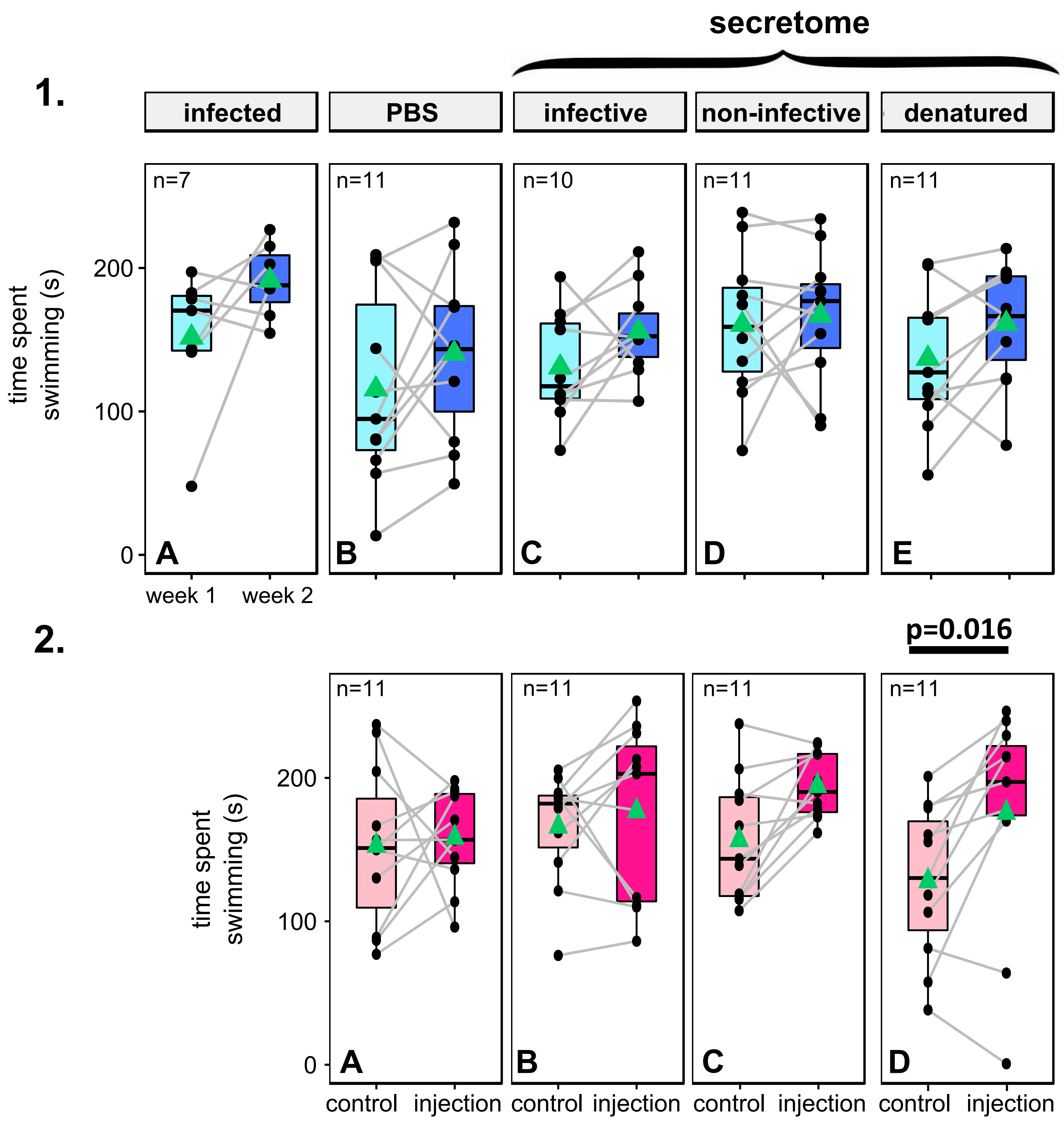

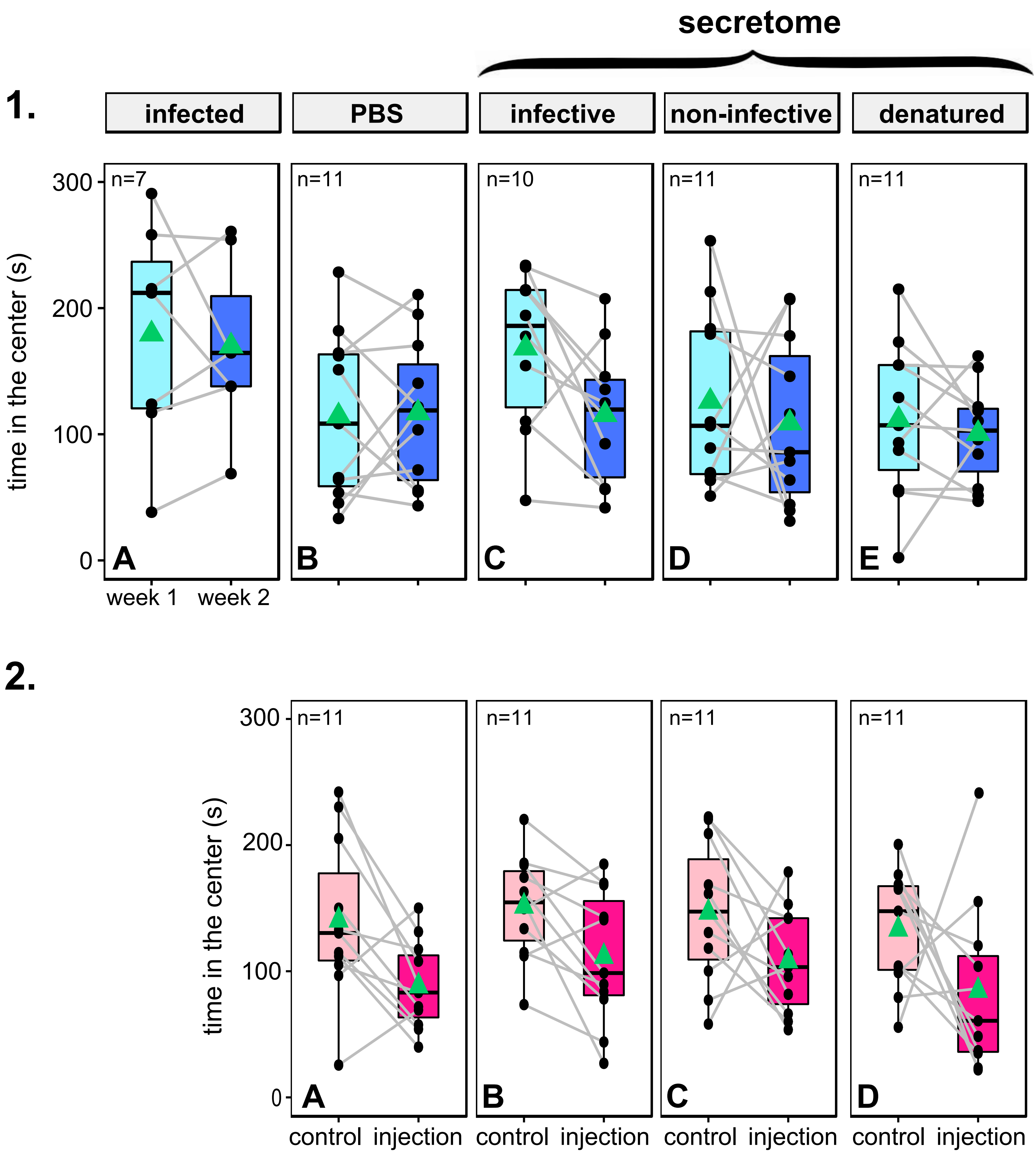

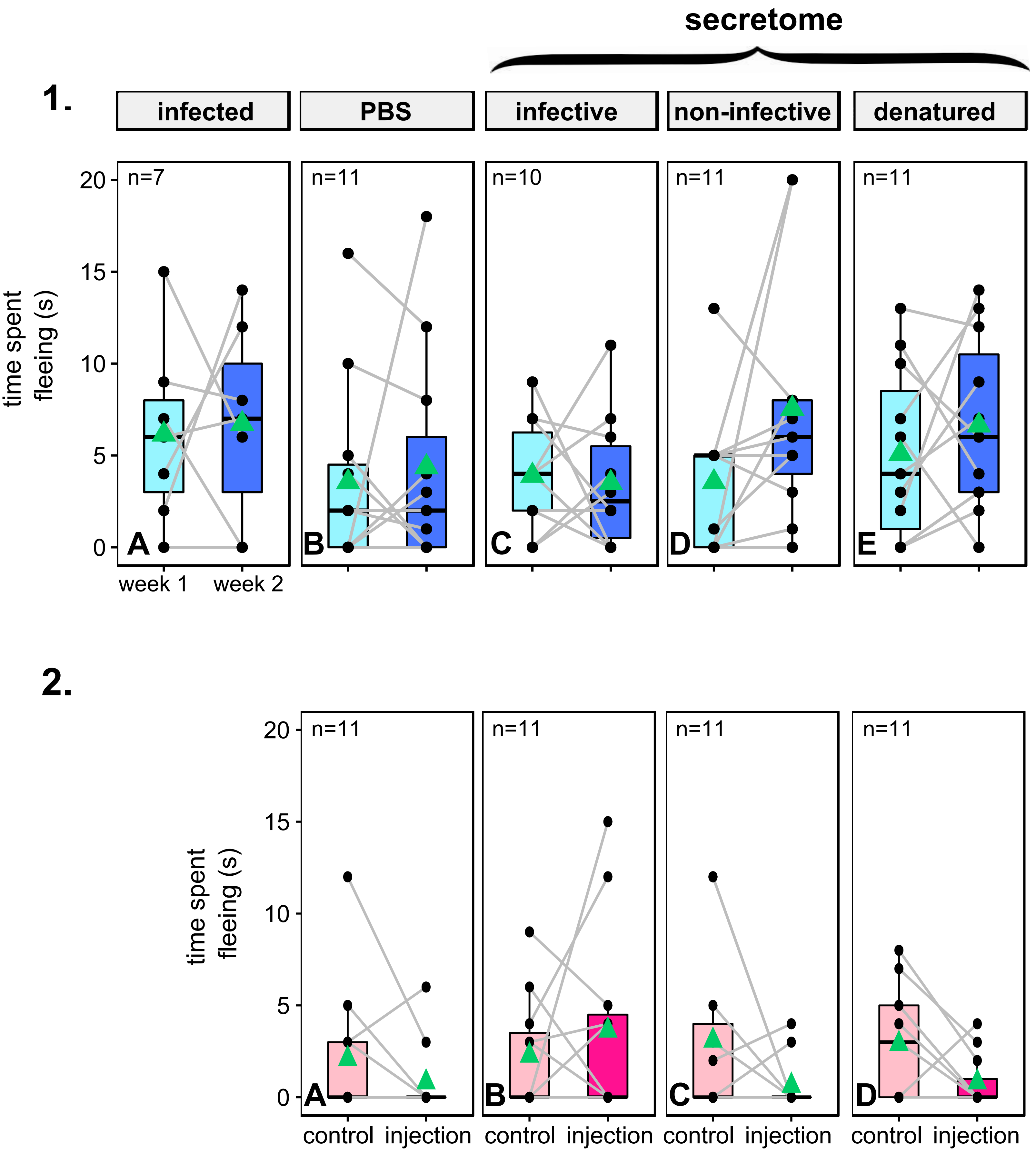

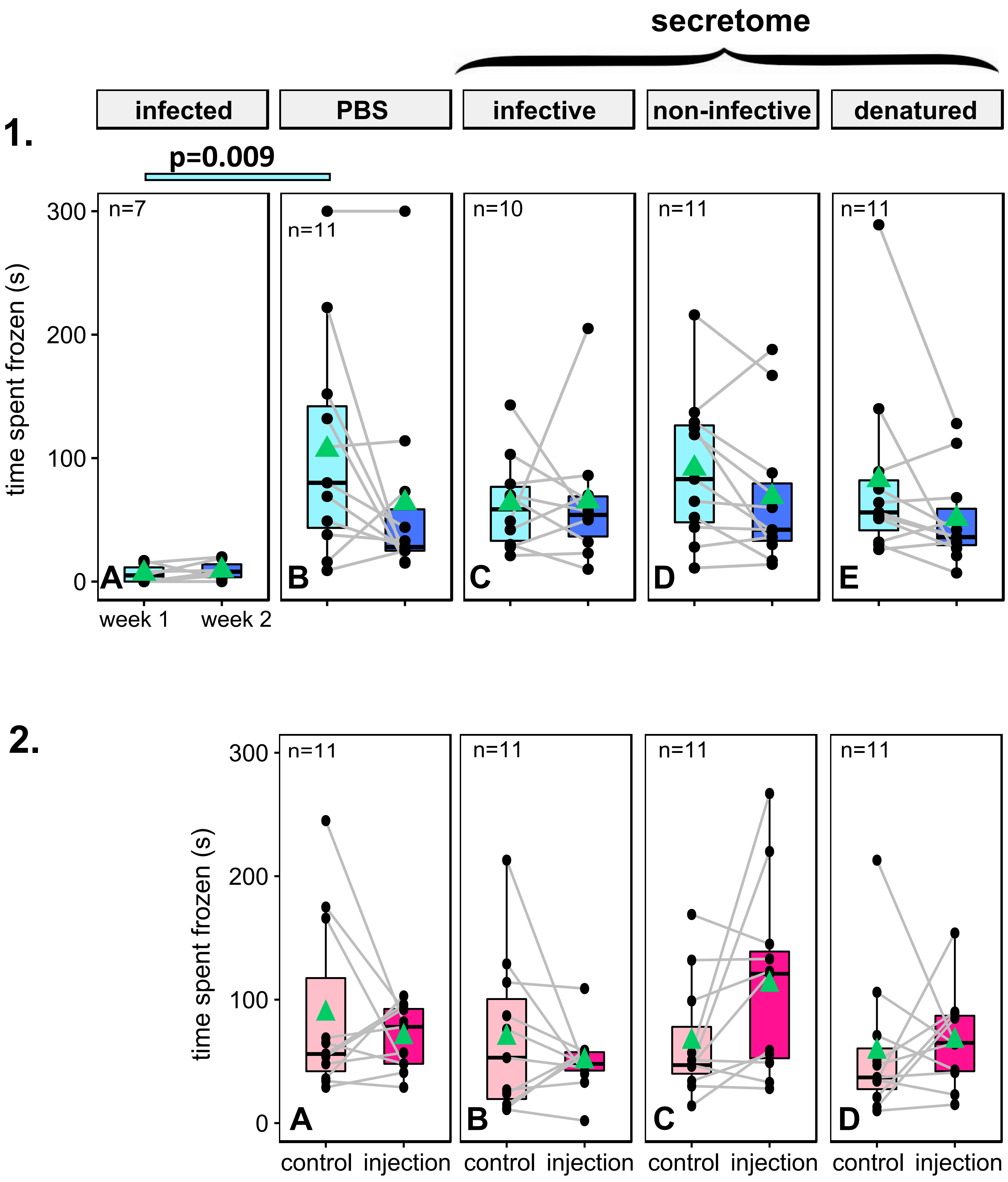

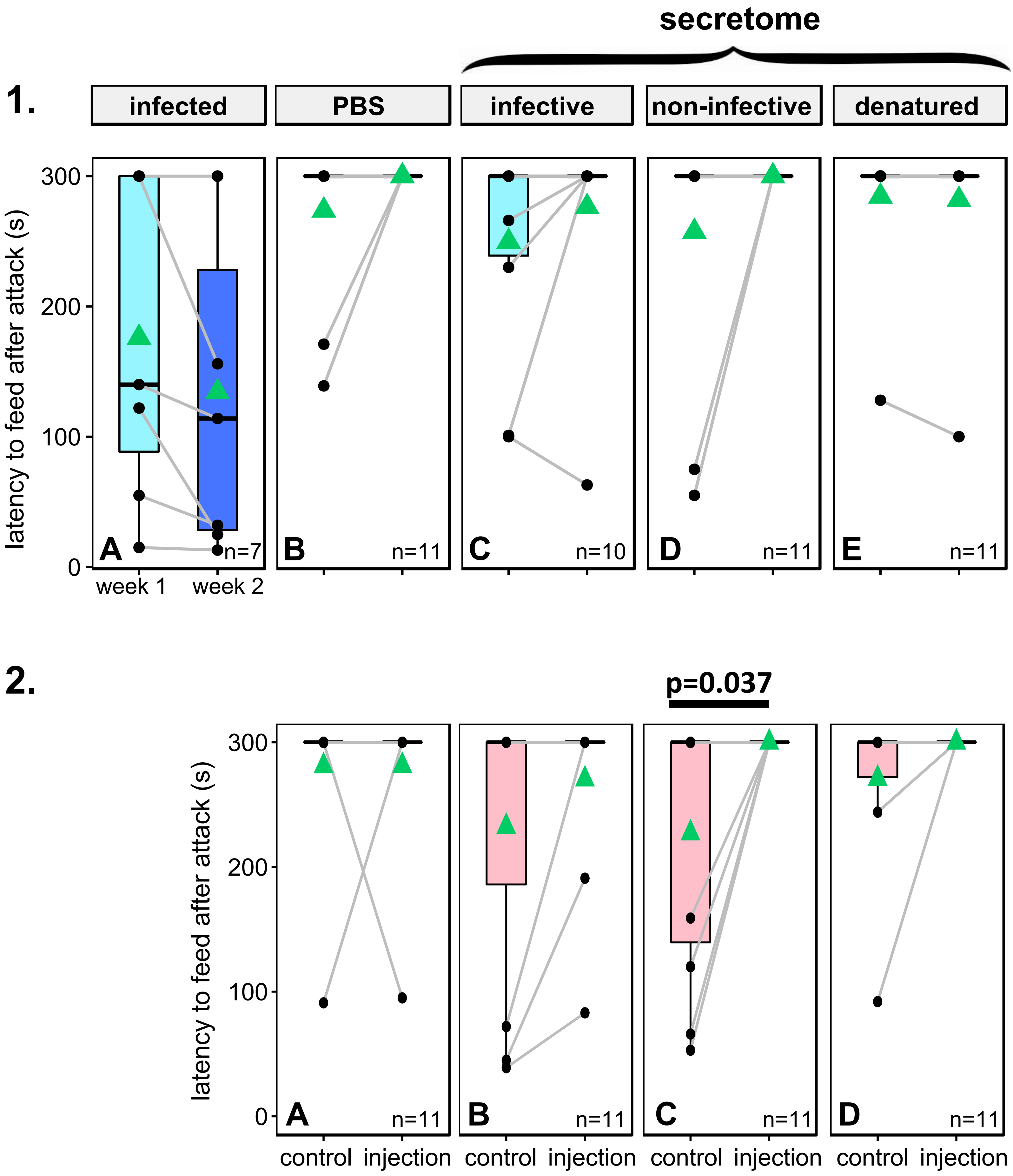
