## Supplementary tables for "The secretome of a parasite alters its host’s behaviour but does not recapitulate the behavioural response to infection"

**Table 1.** Results of the statistical comparisons performed with the lmer function to determine if the secretome injections change the behaviour (latency to leave the acclimation zone) of non-infected fish from the Témiscouata population. Two distinct analyses are reported: the first analysis was performed with reference groups only (PBS and parasitized), and the second was performed with the groups exposed to secretome only. SE=Standard Error.

| Behaviour | Treatments included in the lmer model | Pairwise comparisons | estimate | SE | T ratio | P-value |
|---|---|---|---|---|---|---|
| Exit latency | Témiscouata Reference groups | Control PBS – Injection PBS (effect of injection?) | -0.544 | 0.567 | -0.959 | 0.774 |
| | | Control PBS - Control parasitized (effect of infection ?) | -0.516 | 1.026 | -0.503 | 0.958 |
| | | Control parasitized –Injection parasitized (effect of week ?) | 0.203 | 0.710 | 0.286 | 0.992 |
| | Témiscouata Secretome groups | Control secretome_big – Injection secretome_big | 0.737 | 0.590 | 1.250 | 0.809 |
| | | Control secretome_small – Injection secretome_small | 0.805 | 0.562 | 1.431 | 0.708 |
| | | Control secretome_denatured – Injection secretome_denatured | -0.431 | 0.562 | -0.766 | 0.971 |

**Table 2.** Results of the statistical comparisons performed with the lmer function to determine if the secretome injections change the behaviour (time spent swimming) of non-infected fish from the Témiscouata population. Two distinct analyses are reported: the first analysis was performed with reference groups only (PBS and parasitized), and the second was performed with the groups exposed to secretome only. SE=Standard Error.

| Behaviour | Treatments included in the lmer model | Pairwise comparisons | estimate | SE | T ratio | P-value |
|---|---|---|---|---|---|---|
| Swim | Témiscouata Reference groups | Control PBS – Injection PBS (effect of injection?) | -24.931 | 17.405 | -1.432 | 0.497 |
| | | Control PBS - Control parasitized (effect of infection ?) | 34.954 | 27.790 | 1.258 | 0.596 |
| | | Control parasitized –Injection parasitized (effect of week ?) | -39.714 | 21.818 | -1.820 | 0.297 |
| | Témiscouata Secretome groups | Control secretome_big – Injection secretome_big | -24.960 | 13.374 | -1.866 | 0.440 |
| | | Control secretome_small – Injection secretome_small | -6.315 | 12.751 | -0.495 | 0.996 |
| | | Control secretome_denatured – Injection secretome_denatured | -24.250 | 12.751 | -1.902 | 0.420 |

**Table 3.** Results of the statistical comparisons performed with the lmer function to determine if the secretome injections change the behaviour (center time) of non-infected fish from the Témiscouata population. Two distinct analyses are reported: the first analysis was performed with reference groups only (PBS and parasitized), and the second was performed with the groups exposed to secretome only. SE=Standard Error.

| Behaviour | Treatments included in the lmer model | Pairwise comparisons | estimate | SE | T ratio | P-value |
|---|---|---|---|---|---|---|
| Center time | Témiscouata Reference groups | Control PBS – Injection PBS (effect of injection?) | -2.455 | 17.629 | -0.139 | 0.999 |
| | | Control PBS - Control parasitized (effect of infection ?) | 4.639 | 27.988 | 0.166 | 0.998 |
| | | Control parasitized –Injection parasitized (effect of week ?) | 9.520 | 22.099 | 0.431 | 0.972 |
| | Témiscouata Secretome groups | Control secretome_big – Injection secretome_big | 52.802 | 23.674 | 2.230 | 0.252 |
| | | Control secretome_small – Injection secretome_small | 17.145 | 22.572 | 0.760 | 0.972 |
| | | Control secretome_denatured – Injection secretome_denatured | 11.196 | 22.572 | 0.496 | 0.996 |

**Table 4.** Results of the statistical comparisons performed with the lmer function to determine if the secretome injections change the behaviour (latency to feed before attack) of non-infected fish from the Témiscouata population. Two distinct analyses are reported: the first analysis was performed with reference groups only (PBS and parasitized), and the second was performed with the groups exposed to secretome only. SE=Standard Error.

| **Behaviour** | **Treatments included in the lmer model** | **Pairwise comparisons** | **estimate** | **SE** | **T ratio** | **P-value** |
|---|---|---|---|---|---|---|
| Feed before | Témiscouata Reference groups | Control PBS – Injection PBS (effect of injection?) | -32.182 | 33.998 | -0.947 | 0.781 |
| | | Control PBS - Control parasitized (effect of infection ?) | -135.854 | 49.930 | -2.721 | 0.049* |
| | | Control parasitized –Injection parasitized (effect of week ?) | 52.286 | 42.619 | 1.227 | 0.619 |
| | Témiscouata Secretome groups | Control secretome_big – Injection secretome_big | -31.400 | 42.261 | -0.743 | 0.975 |
| | | Control secretome_small – Injection secretome_small | -46.727 | 40.294 | -1.160 | 0.852 |
| | | Control secretome_denatured – Injection secretome_denatured | -69.545 | 40.294 | -1.726 | 0.526 |

**Table 5.** Results of the statistical comparisons performed with the lmer function to determine if the secretome injections change the behaviour (time spent fleeing after attack) of non-infected fish from the Témiscouata population. Two distinct analyses are reported: the first analysis was performed with reference groups only (PBS and parasitized), and the second was performed with the groups exposed to secretome only. SE=Standard Error.

| Behaviour | Treatments included in the lmer model | Pairwise comparisons | estimate | SE | T ratio | P-value |
|---|---|---|---|---|---|---|
| Flee | Témiscouata Reference groups | Control PBS – Injection PBS (effect of injection?) | -0.223 | 0.506 | -0.441 | 0.971 |
| | | Control PBS - Control parasitized (effect of infection ?) | 0.990 | 0.737 | 1.343 | 0.543 |
| | | Control parasitized –Injection parasitized (effect of week ?) | 0.036 | 0.634 | 0.057 | 1.0 |
| | Témiscouata Secretome groups | Control secretome_big – Injection secretome_big | 0.215 | 0.476 | 0.451 | 0.997 |
| | | Control secretome_small – Injection secretome_small | -1.010 | 0.454 | -2.226 | 0.254 |
| | | Control secretome_denatured – Injection secretome_denatured | -0.496 | 0.454 | -1.093 | 0.880 |

**Table 6.** Results of the statistical comparisons performed with the lmer function to determine if the secretome injections change the behaviour (time spent frozen after attack) of non-infected fish from the Témiscouata population. Two distinct analyses are reported: the first analysis was performed with reference groups only (PBS and parasitized), and the second was performed with the groups exposed to secretome only. SE=Standard Error.

| Behaviour | Treatments included in the lmer model | Pairwise comparisons | estimate | SE | T ratio | P-value |
|---|---|---|---|---|---|---|
| Frozen | Témiscouata Reference groups | Control PBS – Injection PBS (effect of injection?) | 2.428 | 1.081 | 2.247 | 0.148 |
| | | Control PBS - Control parasitized (effect of infection ?) | -6.580 | 1.887 | -3.487 | 0.009* |
| | | Control parasitized –Injection parasitized (effect of week ?) | -0.750 | 1.355 | -0.554 | 0.944 |
| | Témiscouata Secretome groups | Control secretome_big – Injection secretome_big | 0.084 | 0.210 | 0.398 | 0.999 |
| | | Control secretome_small – Injection secretome_small | 0.338 | 0.200 | 1.686 | 0.550 |
| | | Control secretome_denatured – Injection secretome_denatured | 0.479 | 0.200 | 2.390 | 0.190 |

**Table 7.** Results of the statistical comparisons performed with the lmer function to determine if the secretome injections change the behaviour (latency to feed after attack) of non-infected fish from the Témiscouata population. Two distinct analyses are reported: the first analysis was performed with reference groups only (PBS and parasitized), and the second was performed with the groups exposed to secretome only. SE=Standard Error.

| Behaviour | Treatments included in the lmer model | Pairwise comparisons | estimate | SE | T ratio | P-value |
|---|---|---|---|---|---|---|
| Feed after | Témiscouata Reference groups | Control PBS – Injection PBS (effect of injection?) | -26.364 | 16.528 | -1.595 | 0.406 |
| | | Control PBS - Control parasitized (effect of infection ?) | -77.414 | 38.505 | -2.010 | 0.214 |
| | | Control parasitized –Injection parasitized (effect of week ?) | 41.714 | 20.719 | 2.013 | 0.220 |
| | Témiscouata Secretome groups | Control secretome_big – Injection secretome_big | -26.600 | 20.251 | -1.314 | 0.776 |
| | | Control secretome_small – Injection secretome_small | -42.727 | 19.308 | -2.213 | 0.260 |
| | | Control secretome_denatured – Injection secretome_denatured | 2.545 | 19.308 | 0.132 | 1.000 |

**Table 8.** Results of the statistical comparisons performed with the lmer function to determine if the secretome injections change the behaviour (latency to leave the acclimation zone) of non-infected fish from the anadromous population. Two distinct analyses are reported: the first analysis included all the groups including the reference (PBS), and the second analysis was performed with the groups exposed to secretome only. SE=Standard Error.

| Behaviour | Treatments included in the lmer model | Pairwise comparisons | estimate | SE | T ratio | P-value |
|---|---|---|---|---|---|---|
| Exit latency | Anadromous Reference | Control PBS – Injection PBS (effect of injection?) | -1.163 | 1.492 | -0.779 | 0.993 |
| | Anadromous Secretome groups | Control secretome_big – Injection secretome_big | 2.216 | 1.421 | 1.559 | 0.630 |
| | | Control secretome_small – Injection secretome_small | -0.04 | 1.421 | -0.029 | 1.000 |
| | | Control secretome_denatured – Injection secretome_denatured | -2.133 | 1.421 | -1.501 | 0.666 |

**Table 9.** Results of the statistical comparisons performed with the lmer function to determine if the secretome injections change the behaviour (time spent swimming) of non-infected fish from the anadromous population. Two distinct analyses are reported: the first analysis included all the groups including the reference (PBS), and the second analysis was performed with the groups exposed to secretome only. SE=Standard Error.

| Behaviour | Treatments included in the lmer model | Pairwise comparisons | estimate | SE | T ratio | P-value |
|---|---|---|---|---|---|---|
| Swim | Anadromous Reference | Control PBS – Injection PBS (effect of injection?) | -5.652 | 15.214 | -0.371 | 1.000 |
| | Anadromous Secretome groups | Control secretome_big – Injection secretome_big | -10.724 | 13.668 | -0.785 | 0.968 |
| | | Control secretome_small – Injection secretome_small | -37.674 | 13.668 | -2.756 | 0.091 |
| | | Control secretome_denatured – Injection secretome_denatured | -47.641 | 13.668 | -3.486 | 0.016* |

**Table 10.** Results of the statistical comparisons performed with the lmer function to determine if the secretome injections change the behaviour (center time) of non-infected fish from the anadromous population. Two distinct analyses are reported: the first analysis included all the groups including the reference (PBS), and the second analysis was performed with the groups exposed to secretome only. SE=Standard Error.

| Behaviour | Treatments included in the lmer model | Pairwise comparisons | estimate | SE | T ratio | P-value |
|---|---|---|---|---|---|---|
| Center time | Anadromous Reference | Control PBS – Injection PBS (effect of injection?) | 52.125 | 20.850 | 2.500 | 0.209 |
| | Anadromous Secretome groups | Control secretome_big – Injection secretome_big | 39.853 | 20.761 | 1.920 | 0.400 |
| | | Control secretome_small – Injection secretome_small | 38.520 | 20.762 | 1.855 | 0.438 |
| | | Control secretome_denatured – Injection secretome_denatured | 48.573 | 20.762 | 2.340 | 0.193 |

**Table 11.** Results of the statistical comparisons performed with the lmer function to determine if the secretome injections change the behaviour (latency to feed before attack) of non-infected fish from the anadromous population. Two distinct analyses are reported: the first analysis included all the groups including the reference (PBS), and the second analysis was performed with the groups exposed to secretome only. SE=Standard Error.

| Behaviour | Treatments included in the lmer model | Pairwise comparisons | estimate | SE | T ratio | P-value |
|---|---|---|---|---|---|---|
| Feed before | Anadromous Reference | Control PBS – Injection PBS (effect of injection?) | -49.000 | 31.117 | -1.575 | 0.762 |
| | Anadromous Secretome groups | Control secretome_big – Injection secretome_big | -2.183 | 1.418 | -1.539 | 0.642 |
| | | Control secretome_small – Injection secretome_small | -6.862 | 1.418 | -4.839 | 0.0004* |
| | | Control secretome_denatured – Injection secretome_denatured | -4.690 | 1.418 | -3.307 | 0.025* |

**Table 12.** Results of the statistical comparisons performed with the lmer function to determine if the secretome injections change the behaviour (time spent fleeing after attack) of non-infected fish from the anadromous population. Two distinct analyses are reported: the first analysis included all the groups including the reference (PBS), and the second analysis was performed with the groups exposed to secretome only. SE=Standard Error.

| Behaviour | Treatments included in the lmer model | Pairwise comparisons | estimate | SE | T ratio | P-value |
|---|---|---|---|---|---|---|
| Flee | Anadromous Reference | Control PBS – Injection PBS (effect of injection?) | 0.453 | 0.420 | 1.079 | 0.958 |
| | Anadromous Secretome groups | Control secretome_big – Injection secretome_big | -0.247 | 0.458 | -0.528 | 0.995 |
| | | Control secretome_small – Injection secretome_small | 0.780 | 0.458 | 1.704 | 0.539 |
| | | Control secretome_denatured – Injection secretome_denatured | 0.776 | 0.458 | 1.695 | 0.545 |

**Table 13.** Results of the statistical comparisons performed with the lmer function to determine if the secretome injections change the behaviour (time spent frozen after attack) of non-infected fish from the anadromous population. Two distinct analyses are reported: the first analysis included all the groups including the reference (PBS), and the second analysis was performed with the groups exposed to secretome only. SE=Standard Error.

| Behaviour | Treatments included in the lmer model | Pairwise comparisons | estimate | SE | T ratio | P-value |
|---|---|---|---|---|---|---|
| Freeze | Anadromous Reference | Control PBS – Injection PBS (effect of injection?) | 0.061 | 0.256 | 0.236 | 1.000 |
| | Anadromous Secretome groups | Control secretome_big – Injection secretome_big | 0.141 | 0.271 | 0.518 | 0.995 |
| | | Control secretome_small – Injection secretome_small | -0.509 | 0.271 | -1.878 | 0.433 |
| | | Control secretome_denatured – Injection secretome_denatured | -0.331 | 0.271 | -1.220 | 0.824 |

**Table 14.** Results of the statistical comparisons performed with the lmer function to determine if the secretome injections change the behaviour (latency to feed after attack) of non-infected fish from the anadromous population. Two distinct analyses are reported: the first analysis included all the groups including the reference (PBS), and the second analysis was performed with the groups exposed to secretome only. SE=Standard Error.

| Behaviour | Treatments included in the lmer model | Pairwise comparisons | estimate | SE | T ratio | P-value |
|---|---|---|---|---|---|---|
| Feed after | Anadromous Reference | Control PBS – Injection PBS (effect of injection?) | -67.636 | 9405.64 | -0.007 | 1.000 |
| | Anadromous Secretome groups | Control secretome_big – Injection secretome_big | -11330.909 | 9027.457 | -1.255 | 0.806 |
| | | Control secretome_small – Injection secretome_small | -28468.545 | 9027.457 | -3.154 | 0.037 * |
| | | Control secretome_denatured – Injection secretome_denatured | -12951.273 | 9027.457 | -1.435 | 0.706 |
