## Supplementary methods for "The secretome of a parasite alters its host’s behaviour but does not recapitulate the behavioural response to infection"

**Electronic Supplementary Information**

### Collecting secretome from the non-infective and infective phases

Worms were dissected from wild sticklebacks. To wash the worm thoroughly after extraction from the host body cavity, we placed it in a 2mL square well of a 96-well plate filled with 1 mL of PBS (Phosphate Buffered Saline, pH 7.4 Life Technologies) during 1 min. After 1 min, the PBS was removed and replaced by 1 mL of fresh PBS. During washing, the plate was covered with aluminium foil to protect the worm from light, and the plate was not shaken in order to prevent the secretion by the worm of proteins involved in the stress response. After washing, each worm was put in a 2 mL tube of PBS. The tube was covered with aluminium foil to protect the worm from light and placed in a water recipient from the lake for 2 hours. We then removed the worm from the tube.

### Protein content of secretome

Prior to injections, we determined the protein concentration of each secretome mixture using a Pierce Coomassie (Bradford) Protein Assay Kit (Fisher Scientific). Bovine Serum Albumin (BSA) (2 mg/mL) was used as standard. The concentrations obtained for the infective secretome, the non-infective secretome and the denatured secretome were respectively 24.08 µg/mL, 6.47 µg/mL and 40.48 µg/mL. The heat-denatured secretome was used as a control of the intact infective secretome, to test for the role of the activity of the proteins in behavioural alterations. The denatured secretome was diluted to the concentration of the infective secretome in order to remove potential concentration effects, and analyze only the effects of the activity of the proteins on behaviour. On the other hand, the concentration of the non-infective secretome was not adjusted to the concentration of the infective secretome to illustrate secretome concentration variation when worms of different weighs are put in a saline solution of constant volume mimicking the abdominal cavity of the host. We visualized proteins for each secretome mixture on a SDS-PAGE (resolving gel 12%) using a Pierce Silver Stain Kit (Fisher Scientific). Before running a gel, each mixture (100 µL) was precipitated with Trichloroacetic Acid (TCA) to facilitate protein visualization at low concentration. A yeast protein extract was used as a positive control and a Blue pre-stained protein standard (BioLabs) was added on the gel. A negative control including the SDS-PAGE loading buffer and water was used. Migration was performed during 60 min at 175 V followed by silver staining. For the three mixtures, proteins of various sizes were effectively separated on gel (Supp Figure 1).

**Sampling of sympatric and allopatric fish hosts**

We caught sticklebacks from Lake Témiscouata using a seine in August 2016. Fish from the allopatric anadromous Isle-Verte population were sampled in tide pools in July 2016 using hand nets. Fish from both populations were juveniles at the time of capture. We brought them to the “Laboratoire Aquatique de Recherche en Sciences Environnementales et Médicales” at Université Laval. Sticklebacks from both populations were maintained in separate 80 L tanks under a photoperiod of 12 h:12 h and a temperature of 15°C reflecting their natural environment conditions. Salinity was at 28 ppt for allopatric fish. We fed fish daily with brine shrimps. All sticklebacks were mature at the start of the injections.

**Behavioural tests**

At the beginning of the protocol, 51 fish from the Témiscouata population were isolated in 2-L tanks filled with freshwater. They were acclimatized to the tanks during two weeks. Fish were then behaviourally tested (see below). After tests with the Témiscouata population were completed, all the 2-L tanks were washed and 44 fish from the Isle-Verte population were isolated in the same 2-L tanks, now filled with saltwater at 28 ppt. They were also acclimatized to the tanks during two weeks and then behaviourally tested. Acclimatization of the two populations to the same 2-L tanks aimed at reducing any potential tank effects. Each fish was assigned an identification number (ID).

We performed behavioural tests in a test aquarium (30 Width x 30 Depth x 60 Length cm) inside which a white box (25 W x 36 D x 57 L cm) was inserted in order to prevent fish reflections on the glass of the aquarium and to optimize tracking during analyses. We filled the box with 25 cm of water depth and water was changed between each fish. The box was opened on the top to allow recording by a camera. The day before the behavioural test of a fish, we enclosed the 2L individual tank with a small white box (14 W x 17 D x 30 L cm) opened on top, to acclimatize the fish to a white environment. From this point the fish was not fed. The day of the test, at the start of the experiment, we put the fish in the acclimation zone inside the white box of the aquarium for 600 seconds. The acclimation zone, which was delimited by a white wall, represented one third of the white box. After 600 seconds, the wall was removed, and 3 behaviours were measured: exploration, thigmotaxis and boldness. Two behaviours were automatically analysed using a tracking module with the Ethovision software (Ethovision XT 11.5, Noldus Information Technology) (54): time spent swimming, a measure of exploration, and time spent in the centre of the aquarium, a measure of thigmotaxis. All the other behaviours were analysed manually. All the videos were analysed blinded by C.S.B.

<u>Exploration</u>

Exploration of a novel environment was measured using 2 variables: 1) Latency to leave the acclimation zone (sec). As soon as the wall was removed, the fish was given 600 seconds to leave the acclimation zone. If the fish never left the acclimation zone, it was given the worst score for exploration (*i.e.* 600 sec). 2) Time spent swimming (sec). As soon as the fish left the acclimation zone (or after 600 sec), time spent swimming by the fish was recorded for 300 seconds.

Thigmotaxis

Thigmotaxis, also referred to as Wall hugging was measured with 1 variable: time spent in the centre of the aquarium (sec). We quantified thigmotaxis as soon as the fish left the acclimation zone (or after 600 sec) and measured for 300 seconds. The centre of the aquarium was defined as a zone at more than 5 cm of the edge of the white box. Fish spending a high amount of time in the centre of the aquarium has low thigmotaxis.

Boldness

The inclination to take risks was measured using 4 variables: 1) Latency to feed before a predator attack (sec). We deposited brine shrimps with a pipette in front of the fish so that it could see it. The fish was given 300 seconds to feed. 2) Time spent fleeing after a predator attack (sec). 3) Time spent frozen after a predator attack (sec). 4) Latency to feed after a predator attack (sec). For these last three variables, as soon as the fish fed after food deposition, or 300 seconds after food deposition, we simulated a predator attack using a heron head model (beak of the model was going back and forth into the water during 5 seconds near the fish head). Time spent fleeing corresponds to the time during which the fish escapes just after the heron head model was removed. Time spent frozen indicates the time during which the fish stays motionless (*i.e.* does not swim a distance greater than its body length) just after the heron head model was removed. Latency to feed after an attack was also measured just after the heron head model was removed. These three variables were determined for 300 seconds.

Following the behavioural tests, we euthanized fish on day 4 of the injection week with an overdose of MS-222 (75 mg/L) and dissected them to confirm that they were infected by

*S. solidus* or not. Fish sex, size and mass, and *S. solidus* mass and number in each fish were noted.

**Statistical analyses**

We used the package lme4 (56) to create a linear mixed model for each population separately including the week (control or injection), the treatment (PBS, “infective secretome”, “non-infective secretome”, “denatured secretome” or infected) and their interaction (week*treatment), the mass, the total length and the sex of the fish as fixed effects and the ID of the fish as random effect. For each analysis, we checked homoscedasticity by plotting residuals and fitted data. We checked for homogeneity of variances using a Levene Test. We tested normality of residuals using graphical assessment (QQ-plot and normal distribution histogram) and a Shapiro-Wilk test. When the data were not normally distributed, we used the Boxcox test from the package MASS (57) to find the best transformation to fit normality (log, square root, square or no transformation). To obtain p-values, we performed pairwise comparisons between weeks and treatments using the package lmerTest. The Tukey method was used to correct for multiple testing and type 1 errors (false discoveries) (58). To confirm our results, we performed additional statistical analyses to validate that our model was not overfitted and that the obtained p-values were accurate. We found that mass of fish was correlated with behaviour, but p-values were not changed when removing this effect from the model.

In the sympatric Témiscouata population, we performed a first analysis with two reference treatments (infected and PBS) to confirm that there was no difference in behaviour before (PBS-control week) and after (PBS-injection week) control injections, and to determine if infected fish (infected-control week) and non-infected fish (PBS-control week) showed significantly different behaviours as expected. The second analysis aimed to test our main predictions and was performed with the secretome treatments only (“infective secretome”, ‘non-infective secretome” and “denatured secretome”) to determine if the secretome injections were able to reproduce the behaviours of infected fish, using each individual as their own control.

In the allopatric anadromous population, fish were not infected by *S. solidus*. The first analysis was thereby performed with all reference (PBS) and secretome treatments, to test if PBS injections changed behaviours between week one and two. To test our predictions about the effects of secretome on behaviour, we performed a second analysis using fish from the three secretome treatments (“infective secretome”, “non-infective secretome” and “denatured secretome”), using each individual as their own control.
